# High-resolution line-scanning reveals distinct visual response properties across human cortical layers

**DOI:** 10.1101/2020.06.30.179762

**Authors:** Andrew T. Morgan, Nils Nothnagel, Lucy. S. Petro, Jozien Goense, Lars Muckli

## Abstract

Our understanding of the human brain relies on advancing noninvasive brain imaging approaches. Characterization of the function of brain circuitry depends on the spatiotemporal correspondence at which recorded signals can be mapped onto underlying neuronal structures and processes. Here we aimed to address key first-stage questions of feasibility, reliability, and utility of line-scanning fMRI as a next generation non-invasive imaging method for human neuroscience research at the mesoscopic scale. Line-scanning can achieve high spatial resolution by employing anisotropic voxels aligned to cortical layers. The method can simultaneously achieve high temporal resolution by limiting acquisition to a very small patch of cortex which is repeatedly acquired as a single frequency-encoded k-space line. We developed multi-echo line-scanning procedures to record cortical layers in humans at high spatial (200 μm) and temporal resolution (100 ms) using ultra high-field 7T fMRI. Quantitative mapping allowed us to identify cortical layers in primary visual cortex (V1) and record functional signals from them while participants viewed movie clips. Analysis of these recordings revealed layer-specific V1 spatial and orientation tuning properties analogous to those previously observed in electrophysiological recordings of non-human primates. We have consequently demonstrated that line-scanning is a powerful non-invasive imaging technique for investigating mesoscopic functional circuits in human cortex.

**Graphical Abstract:** 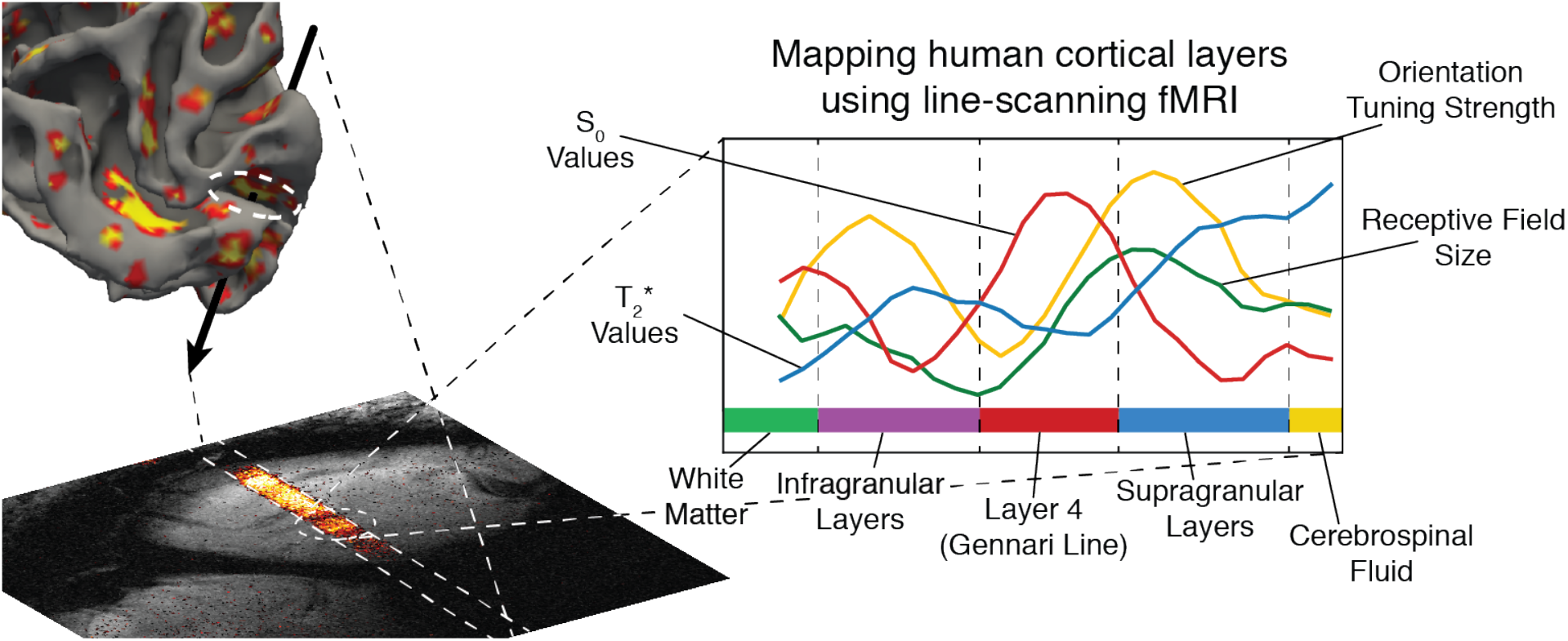

## 1. Introduction

Our understanding of the human brain relies on advancing noninvasive brain imaging approaches. The characterization of brain circuitry and activation depends on the spatiotemporal correspondence in which recorded signals can be mapped to underlying neuronal structures and processes. Ultra high-field (UHF) functional magnetic resonance imaging (fMRI) has advanced the spatial resolution of fMRI to the sub-millimeter range, with 7 Tesla fMRI studies routinely achieving 0.7-0.8 mm^3^ isotropic resolution. This improved resolution enables the discrimination of mesoscopic signals from cortical columns (Martino et al., 2015; Schneider et al., 2019), the identification of laminar fMRI response properties within cortical areas (Kok et al., 2016; Muckli et al., 2015), and lamina-specific mapping of cortical connectivity (Huber et al., 2020).

Promisingly, UHF fMRI continues to augment the spatial scale at which we can access the human brain. A particularly exciting prospect is that UHF fMRI can complement electrophysiology to produce an accurate picture of laminar differences in neural processing (Self et al., 2019). Nevertheless, a number of challenges exist, including the tendency for typical fMRI voxels in humans to be larger than the thickness of individual layers, leading to partial volume effects in which voxels contain an aggregation of signals from multiple layers. Human layer-specific measures must consequently be interpreted in the context of a degree of distortion, where functional “layers” do not map directly onto equivalent anatomical laminae.

While limiting factors such as gradient strength and slew rate affect the achievable spatial resolution of fMRI (Goense et al., 2016), experimenters encounter a general tradeoff between spatial and temporal resolution. One solution to minimize partial volume contamination between layers has been to record anisotropic data-increasing the spatial resolution perpendicular to layers while forfeiting resolution in other directions (Kashyap et al., 2018). This increases the experimenter’s ability to delineate cortical layers while keeping temporal resolution comparable to typical fMRI studies. However, the human brain is highly gyrified, making it difficult to align more than a small portion of the image with cortical layers. Temporal resolution could therefore be further improved by limiting acquisition to a small cortical patch of interest.

In the current work, we aimed to close the gap between layer-dependent fMRI in humans and laminar electrophysiological recordings. Recent studies using fMRI in rodents (Albers et al., 2018; Yu et al., 2014) have shown that line-scanning can achieve high spatial resolution through cortical layers by employing anisotropic data acquisition and can achieve high temporal resolution by limiting acquisition to a very small patch of cortex imaged using a single frequency-encoded k-space line. Motivated by these findings, we developed a line-scanning fMRI procedure to record cortical layers in humans at high spatial (200 μm) and temporal resolution (100 ms) using fMRI. Our approach has a comparable temporal resolution to electrophysiological responses such as event-related potentials or post-stimulus time histograms while preserving a spatial resolution that allows us to resolve cortical layers in human cortex.

We recorded functional signals from cortical layers of primary visual cortex (V1) while participants viewed movie clips. Analysis of these recordings reproduced layer-specific V1 tuning properties previously observed in electrophysiological recordings of non-human primates (Gilbert, 1977; Hubel and Wiesel, 1977; Self et al., 2013). These results therefore demonstrate that line-scanning is a promising and powerful non-invasive imaging technique for investigating local functional circuits in human cortex. Further, this approach lays a precedent for the capability to record sensory or cognitive events at a functionally relevant time scale and at a spatial resolution of mesoscopic cortical circuitry.

## 2. Materials and Methods

### Subjects

Three healthy individuals (2 female; ages = 21, 28, 33) with normal or corrected-to-normal vision gave written informed consent to participate in this study, in accordance with the institutional guidelines of the local ethics committee of the College of Medical, Veterinary and Life Sciences at the University of Glasgow (Project no. 302122).

### Experimental Design

Participants viewed four movie clips previously used in the Human Connectome Project (Cutting et al., 2012). Clips came from the HO (Hollywood movie) set, including ‘Inception,’ ‘The Social Network,’ ‘Oceans 11,’ and ‘Vimeo repeat’ clips [266.8, 298.6, 289.3, and 123.4 seconds, respectively]. Movie clips spanned 9.47° × 16.83° visual angle. Subjects passively viewed each clip while maintaining fixation by focusing on a central red fixation cross. Clips were shown to participants during their first scanning session and were repeated in a second session.

### Comparison of saturation strategies

The first methodological challenge to address was to achieve line-scanning under restrictive radiofrequency (RF) power limitations (specific absorption rate: SAR). We implemented a line-scanning sequence by adding regional saturation bands to a 2D acquisition slice and compared several strategies to choose our final protocol (Figure 1). The initial strategy was to add two regional saturation bands of 40 mm width and an effective gap of 3 mm in the slice center. We tested two saturation pulses offered from the vendor: ‘regular’ with moderate edge sharpness (Figure 1A), and ‘asymmetric’ with one sharp edge (Figure 1B). The second strategy was a scheme of multiple gradient-spoiled Shinnar-LeRoux (SLR) saturation pulses generated using the Matpulse package (Matson, 1994) to further increase line sharpness. Adjacent to the slice center, we also placed two 40 mm saturation bands with a gap of 10 mm and moderate edge sharpness, and two 10 mm saturation bands with a gap of 3 mm and sharp edges (Figure 1C; SLR parameters: maximum-phase, duration 2.2/7.1 ms, passband ripple 0.1%, stopband ripple 0.1%, bandwidth 2200/4400 Hz). Flip angles for the saturation bands were chosen to be 90° or as high as SAR limitations of the scanner would permit (minimal flip angle of 65° for the inner band and 80° for the outer band). We compared SAR levels and RF peak power for all approaches as reported by the safety monitor of our device (Table 1).

**Figure 1.**
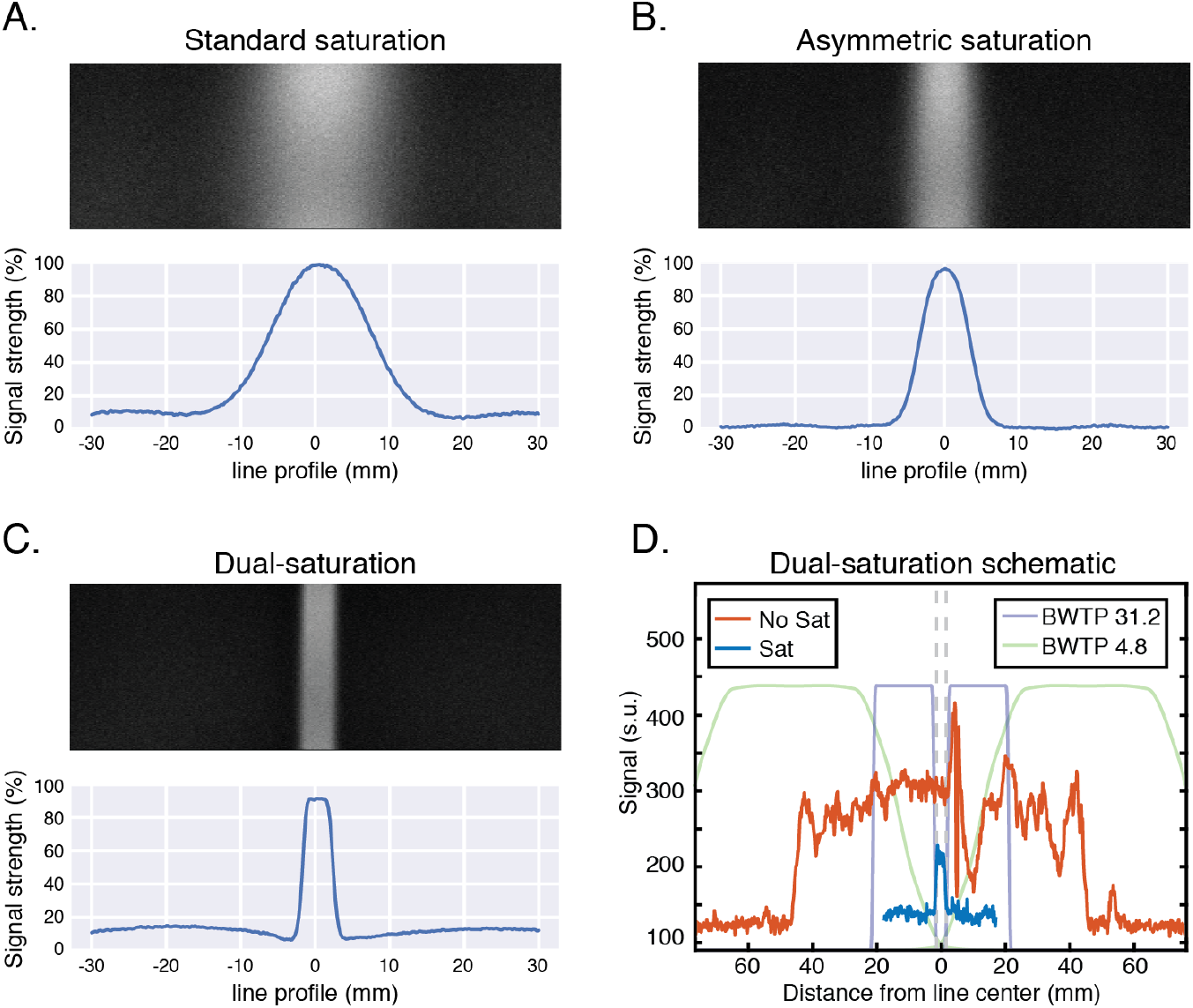
Profiles from possible line-scanning protocols. A. Regular saturation regions with moderate edge sharpness. B. Siemens’ asymmetric saturation regions with sharper edges. C. Customized saturation regions, using a train of four low-energy SLR saturation pulses. D. Schematic illustration of the placement of saturation regions in the dual-saturation scheme and their effect on signal recorded *in vivo*.

**Table 1.**
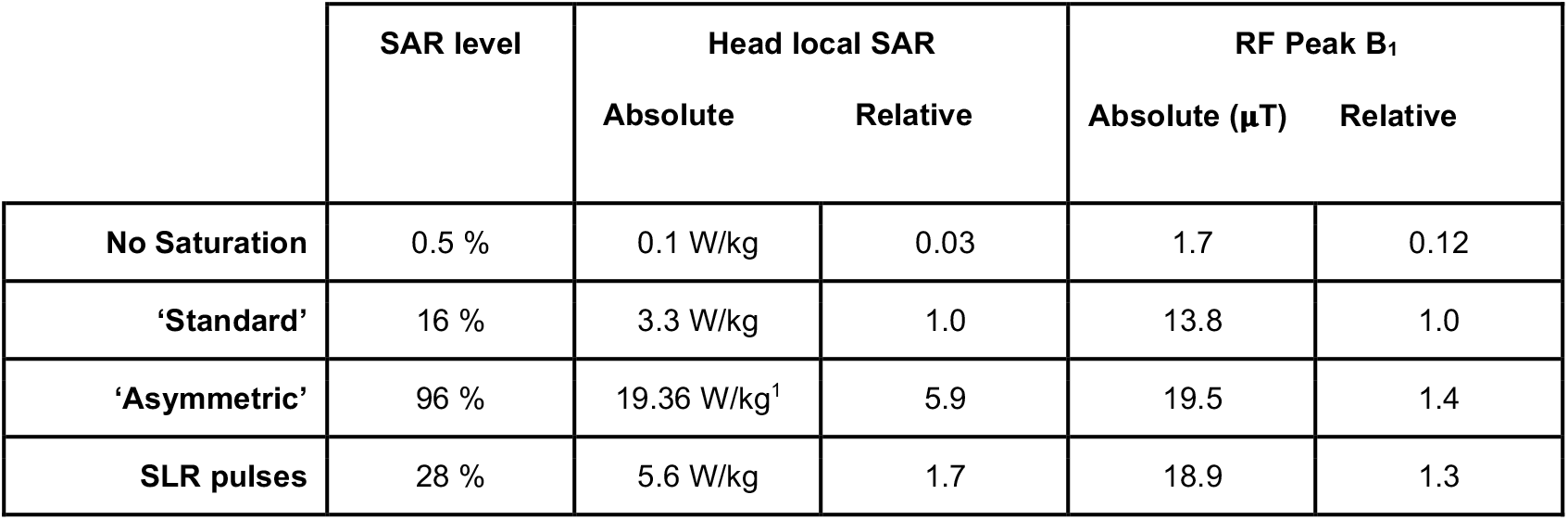
SAR levels and RF power comparison for line-scanning strategies. Values taken from the RF Safety Watchdog of the Siemens Magnetom Terra system. Relative values are reported as divided by the values for the ‘standard’ regional saturation pulses.

|  | SAR level | Head local SAR |  | RF Peak B<sub>1</sub> |  |
| --- | --- | --- | --- | --- | --- |
| | | Absolute | Relative | Absolute (μT) | Relative |
| **No Saturation** | 0.5 % | 0.1 W/kg | 0.03 | 1.7 | 0.12 |
| **'Standard'** | 16 % | 3.3 W/kg | 1.0 | 13.8 | 1.0 |
| **'Asymmetric'** | 96 % | 19.36 W/kg<sup>1</sup> | 5.9 | 19.5 | 1.4 |
| **SLR pulses** | 28 % | 5.6 W/kg | 1.7 | 18.9 | 1.3 |

### Line-scan session planning

A second methodological challenge was planning around cortical folding. We scrutinized cortical folding patterns in individual subjects prior to acquisition to find flat portions of cortex in the area of interest (foveal or parafoveal V1). This minimized partial volume effects caused by cortical layers curving through anisotropic voxels. We generated 3D cortical white matter surface models for each subject using the Freesurfer package (Dale et al., 1999). We then marked vertices with the lowest 10% of both Gaussian and mean curvature values on each surface, as these geometric surface metrics, if both of their values are close to zero, indicate surface flatness. For our experiment, we chose flat cortical patches in foveal or parafoveal V1 at least 3 mm wide where the line-scanning frequency encoding direction would match the vertex normal of this patch. Additionally, to make sure we could adequately saturate signals originating outside the region of interest (ROI), we rotated the imaging plane around the vertex normal and calculated the amount of signal in the imaging plane at each rotation angle that fell outside the saturation area (farther than 73 mm from the vertex). We subsequently selected the angle for the imaging plane that minimized this undesired signal. During recruitment and planning, participants were not scanned if they did not have an imaging plane with less than 95% of total signal falling inside the saturated area.

### fMRI acquisition

Data were collected using a 7T Siemens Magnetom Terra system (Siemens Healthcare, Erlangen, Germany) equipped with a 1Tx/32Rx-channel head coil (Nova Medical Inc., MA, USA) and an SC72 gradient. The head coil was equipped with a rear-facing mirror through which participants viewed the movie clips via a PROPixx projector system (VPixx Technologies, Saint-Bruno, QC, Canada). Participants’ heads were tilted (positive rotation around x-axis of head; chin up) to allow them to view the entire projector screen. We placed high-permittivity pads (calcium titanate in deuterated water; produced by Leiden University) directly on the back of the head centered on the visual cortex to locally increase B_1_ (Brink et al., 2014; Teeuwisse et al., 2012).

At the beginning of each session, we recorded a 3D localizer for alignment purposes (TR=37 ms, TE=10.1 ms, FOV=180×180×30 mm^3^, matrix=180×180×30, flip angle=27°, readout BW=60 Hz/px, GRAPPA=3, Partial Fourier=7/8 in slice and phase directions). Once recorded, this localizer was exported to an external computer and aligned to a previously recorded T_1_-weighted image used to create the 3D surface models used during planning. Alignment was performed semi-automatically using the ITK-SNAP software (Yushkevich et al., 2006). This procedure allowed us to translate the pre-defined position and direction of the cortical patch of interest for line-scanning to the current scanning session. We then converted this new position and direction to the format required by Siemens software to define our imaging slice.

Our line-scanning sequence was based on a multi-echo gradient-spoiled 2D-FLASH sequence (TR=100 ms, TEs=12.6, 24.8, 37.0, 49.2 ms [echo-spacing: 12.2 ms; bipolar readout], FOV=51×51×3 mm^3^, matrix=256×256, resolution=0.2×3.0×3.0 mm^3^, flip angle=18°, readout BW=90 Hz/px). Two versions of the line-scanning sequence were used. The first was a profile version, which employed a standard 2D-FLASH phase encoding scheme. This sequence was used with and without saturation to verify the location of the line. The second sequence version deactivated the phase encoding gradient to achieve spatial encoding purely along frequency encoding direction and was used to record functional movie runs. Each scanning session lasted approximately 50 minutes including setup (Scans: Localizers, B_1_ mapping, Profile image [saturated], Profile image [unsaturated], Functional movies [4 runs]).

### Data Reconstruction

Raw data files were exported and converted to ISMRMRD format (Inati et al., 2017). Since line-scan functional runs used bipolar readouts for the 4 echoes, we needed to correct phase differences between odd and even echoes. This was accomplished by adding a phase offset to the data in the frequency domain before the 1D Fourier transform. We optimized phase offset values to maximize correlation between the ten voxels with the highest signal intensity (line peaks) in the last two echoes of each scan.

We performed motion correction in a similar manner, but instead of comparing peak locations, we used the signal around our gray matter ROI (within 15 mm) as our navigator. Since line-scan data are one-dimensional, motion correction could only account for one degree of freedom: translation in the direction of the frequency encoding gradient. By performing motion correction while data were still in the complex domain, we avoided interpolation of data after reconstruction.

We recombined data from different receiver channels based on each channel’s sensitivity to our very small ROI. This effectively discarded data from approximately 26 of the 32 channels. We first defined an ROI (centered on the excited line) in a 2D profile version of our scan. Next, we linearly combined channels using partial least squares regression to emphasize signal from the line (within the ROI) while simultaneously minimizing signal from non-relevant areas. We used partial least squares regression because it effectively creates virtual channels (Blaimer et al., 2009) in the regression process. These channel weights were then used to reconstruct all 2D anatomical profiles and functional line-scan data from the session.

### Anatomical analysis

We labeled gray matter (GM) voxels by manually inspecting 2D profile images to identify the gray matter/white matter boundary and the gray matter/cerebrospinal fluid boundary. Next, we calculated quantitative T_2_* and S_0_ values for each voxel. Using the first 10 volumes of the line-scan functional runs (where signal intensity is highest), we performed log-linear regression relating signal decay across echoes to echo time. We then identified Layer 4 from reduced T_2_* decay times of voxels in middle cortical layers that highlights the Gennari Line (Barbier et al., 2002; Goense et al., 2007; Walters et al., 2003). We labeled gray matter voxels outside Layer 4 as either supragranular or infragranular layers (above or below Layer 4, respectively).

### Functional preprocessing

We preprocessed functional data using the following steps: 1) Time courses from different echoes were optimally combined by weighted summation by a factor *w*, defined by the following equation:

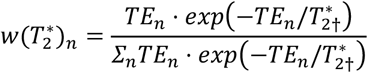

where *TE*_*n*_ is the echo time for a time course and 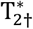 is the baseline T_2_* value in each voxel, as calculated in our anatomical analysis. This combination maximizes sensitivity to the BOLD effect in the functional signal (Kundu et al., 2012). 2) We discarded the first 80 volumes (before steady-state magnetization had been established) and removed polynomial drifts up to 8 degrees. 3) Lastly, we temporally filtered each time course using a Gaussian filter (**σ**=1 s).

### Stimulus preprocessing

Functional analyses were based on an edge energy model applied to the movie clips. We converted movie frames to grayscale and calculated edge energy for 8 orientations (0°, 22.5°, 45°, 67.5°, 90°, 112.5°, 135°, 157.5°) and 4 spatial frequencies (3.7, 2.0, 1.1, 0.6 cycles/degree visual angle), as implemented by default in the GIST model (Oliva and Torralba, 2001). This produced 32 feature time courses for each movie pixel, which were then downsampled to the 10 Hz sampling rate of our line-scan recordings.

### >Receptive field identification

Functional time courses from gray matter voxels were initially averaged to identify the center location of the population receptive field (pRF; Dumoulin and Wandell, 2008) for each subject and session’s cortical patch.

To reduce the dimensionality of this model, we summed across orientation and spatial frequency channels, thus reducing the model to total edge energy at every pixel for every time point of the movie clips. Pixel time courses were convolved with a canonical double-Gamma hemodynamic response function (HRF) and regressed onto each line-scan time course using Fractional Ridge Regression (Rokem and Kay, 2020). We chose the pRF center for further analyses as the peak pixel response when the ratio between the L2-norms of regularized and unregularized coefficients was very low (less than 0.05).

We determined the pRF size for each cortical patch using the following steps: 1) We computed the dot product between each of the 32 (8 orientation × 4 spatial frequency channels) model time courses [dimensions: time × pixels] and a 2D isotropic Gaussian pRF function centered on the previously calculated pRF center location [pixels × 1]. This resulted in time course models of dimensions [time × 32 channels]. 2) We fit the resulting model to the cortical patch time course using Tikhonov regression with a first derivative smoothing regularizer (Aster et al., 2019) that treated spatial frequency and orientation as separate dimensions. 3) The strength of the regularization and the pRF size (30 evenly spaced pRF sizes between 0.1° and 3.5° VA) were treated as hyperparameters that were tuned via cross-validation (Crosse et al., 2016).

### Layer-specific analyses

We repeated the pRF size analysis on a voxel-wise basis (the pRF center kept constant for all gray matter voxels in a cortical patch). Once model features were regressed onto layer-specific voxel activity, we additionally defined orientation tuning sharpness as the difference in response magnitude between preferred (the highest average orientation response for the model, fit to the average of all gray matter time courses) and perpendicular orientations. Since the magnitude of these response difference values differed slightly between subjects and sessions, we normalized tuning strength values across cortical layers (values were scaled so that s.d.=1).

When plotting multiple subject results for cortical depth (pRF size and orientation tuning strength), each subject’s line spanned a normalized cortical thickness (the average cortical thickness between subjects). Since individual subject voxels were therefore not at the same depth, average results were computed using a weighted moving average with a Gaussian kernel (step size = 100 μm, sigma = 100 μm). Additionally, we quantified increases in pRF size and orientation tuning strength by linear regression fit on data points from all subjects between the middle of Layer 4 and the white matter boundary for infragranular layers and between the middle of Layer 4 and the cortical surface for supragranular layers (reported as change in pRF size or orientation tuning strength per 100 μm).

## 3. Results

### Saturation schemes for line-scanning

We first tested a number of saturation schemes to suppress unwanted signal from outside the line-scan area of interest. Our aim was to find a scheme that left the signal from the center of the 3 mm line unchanged, while adequately saturating signal arising from outside that region. Achieving this aim would mean that we could isolate and record from small patches of cortex during functional imaging experiments. Initially, we tested regular regional saturation bands that were placed on either side of the region of interest. This method produced good signal saturation but did not produce sufficiently sharp profiles (Figure 1A). Next, we changed the pulse type of these saturation bands, which improved both profile definition and saturation efficiency (Figure 1B). However, these pulses increased the SAR by a factor of six (see Table 1). The safety monitor on our scanner would therefore not allow a line scanning protocol with a TR lower than 300 ms if these pulses were used.

Therefore, we developed an approach using multiple saturation bands to balance the following requirements: satisfactory suppression of signal from outside the line, sharp profile definition, and reduction of applied RF energy. This approach applied two different saturation pulses on each side of the region of interest. The first pulse was placed at the edge of the region of interest and had a higher bandwidth time product (BWTP; to produce a sharp profile), but slightly lower flip angles (65°). The second pulse was used to widely saturate the signal, so it was placed farther away from the region of interest, had a lower BWTP (which results in a more dispersed saturation profile), and was applied with a higher flip angle (85°). This saturation scheme had slightly reduced saturation efficiency, but showed a much sharper profile, which was achieved with less than a two-fold SAR increase over the regular saturation scheme (Figure 1C; Table 1).

### Cortical layers are discernible from line-scan data

Planning a line-scan experiment requires precise placement of the target volume/line based on anatomical features. Line-scanning can achieve very high resolution in one dimension to examine functional activation of cortical layers. However, the anisotropic voxels recorded in a line-scanning experiment have lower resolution in the other two spatial dimensions. If the high-resolution dimension is perpendicularly aligned to cortical layers, the method can provide a detailed view of hemodynamic processes occurring in cortical layers (Yu et al., 2014). Misalignment would however result in partial-volume effects that would decrease our ability to observe this layer-specific activity.

Figure 2 displays the translation from experimental planning to acquisition of cortical layer data for an individual subject. The gyrencephalic human cortex makes alignment challenging, so we developed a planning procedure to achieve precise alignment in our experiments. We scrutinized cortical folding patterns to find flat portions of cortex (Figure 2A) where the partial volume effects caused by cortical curvature could be avoided. For the current experiment, we chose flat cortical patches in foveal or parafoveal V1 at least 3 mm wide. During each scanning session, we translated the chosen cortical patches to coordinates for the current session and collected 2D anatomical profiles of the line (with and without saturation) to confirm proper placement (Figure 2B). We localized the gray matter for the cortical patch of interest from these profile images and were then able to map infragranular, supragranular and Layer 4 (via the Gennari Line) based on quantitative T_2_* and S_0_ mapping of each voxel (Figure 2C).

**Figure 2.**
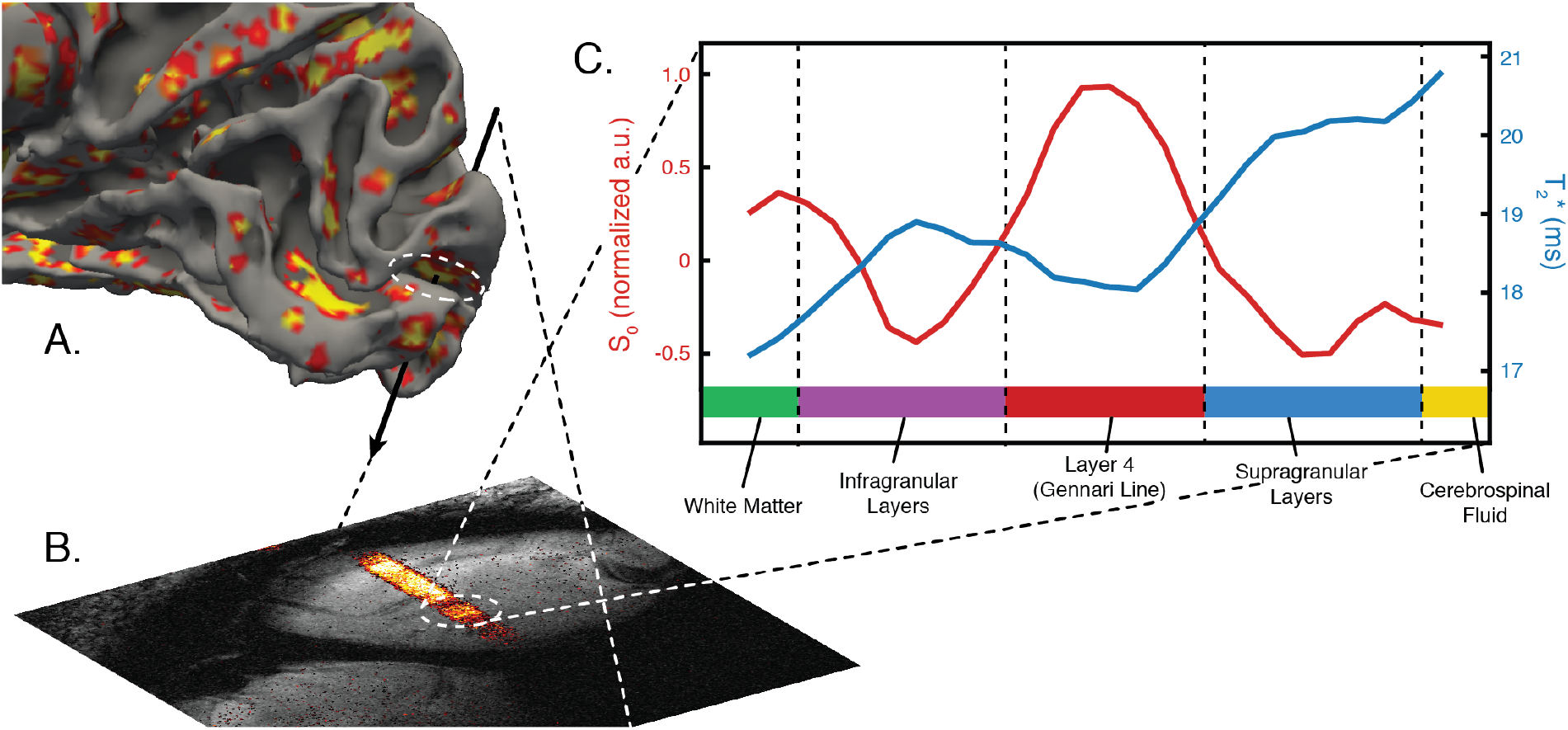
Experimental planning procedures and quantitative mapping of cortical layers. A. A 3D cortical surface model was used to determine the location and angle of the line. Vertices with low curvature values are shown in red and yellow, representing potential locations for line placement. B. A profile scan was performed to confirm correct line placement. An unsaturated profile is shown in grayscale as a background image and the line is overlaid in red/yellow. C. Quantitative mapping was performed (4 echoes) to determine T_2_* and S_0_ values (average values across subjects and sessions are shown). These values were used to determine the boundaries of cortical layers (Layer 4/Line of Gennari, infragranular and supragranular layers).

We were able to consistently map the gray matter patch of interest and cortical layers in each subject and session, as shown in Figure 3. Layer 4 was discernible from reduced T_2_* decay times of voxels in middle cortical layers (Goense et al., 2007; Walters et al., 2003). This shorter T_2_* decay time is due to the Gennari Line, a band of myelinated axons running parallel to the surface of the cortex which corresponds with cellular Layer 4 in primary visual cortex (Fukunaga et al., 2010; Weber et al., 2008).

**Figure 3.**
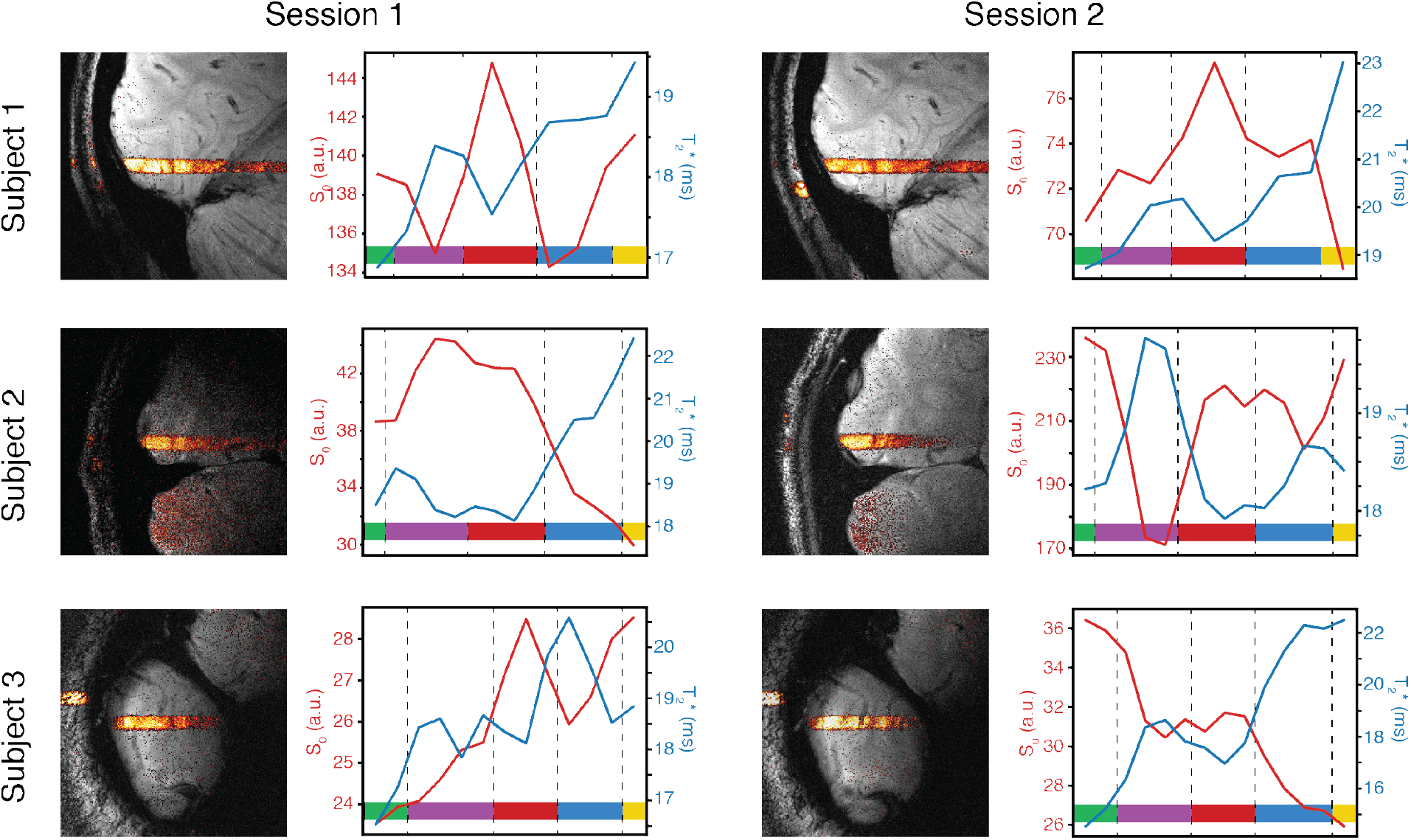
Individual subject quantitative mapping of gray matter. Line-scan profiles and quantitative maps are shown for individual subjects and sessions. Cortical layers/tissues were deduced from a T_2_* peak and double S0 peaks. Colors are the same as in Figure 2: green indicates white matter voxels; purple represents infragranular layers; red represents Layer 4 (Gennari Line); blue represents supragranular layers and yellow represents cerebrospinal fluid.

### Consistent functional responses across sessions

Next, we tested whether cortical patches recorded in each session responded similarly to visual stimulation, as this is an important criterion for the technique being used in neuroscientific experiments. We mapped population receptive fields (Dumoulin and Wandell, 2008) and orientation tuning for each gray matter patch. For each subject, individual session pRFs had similar locations and sizes, with no subject’s pRF differing by more than 1° of visual angle between sessions (center plot of Figure 4). Additionally, orientation tuning was similar across sessions, shown in the surrounding plots of Figure 4. Subject 1’s cortical patch preferred multiple orientations at low frequencies (the center of each plot shows responses to high spatial frequencies and the outside shows responses to low frequencies). Meanwhile, Subjects 2 and 3 had stronger preferences for a single orientation (near vertical and near horizontal, respectively). Taken together with our placement and quantitative mapping results, the consistency of each cortical patch’s functional response across sessions indicates that our procedures are feasible for human neuroscientific studies.

**Figure 4.**
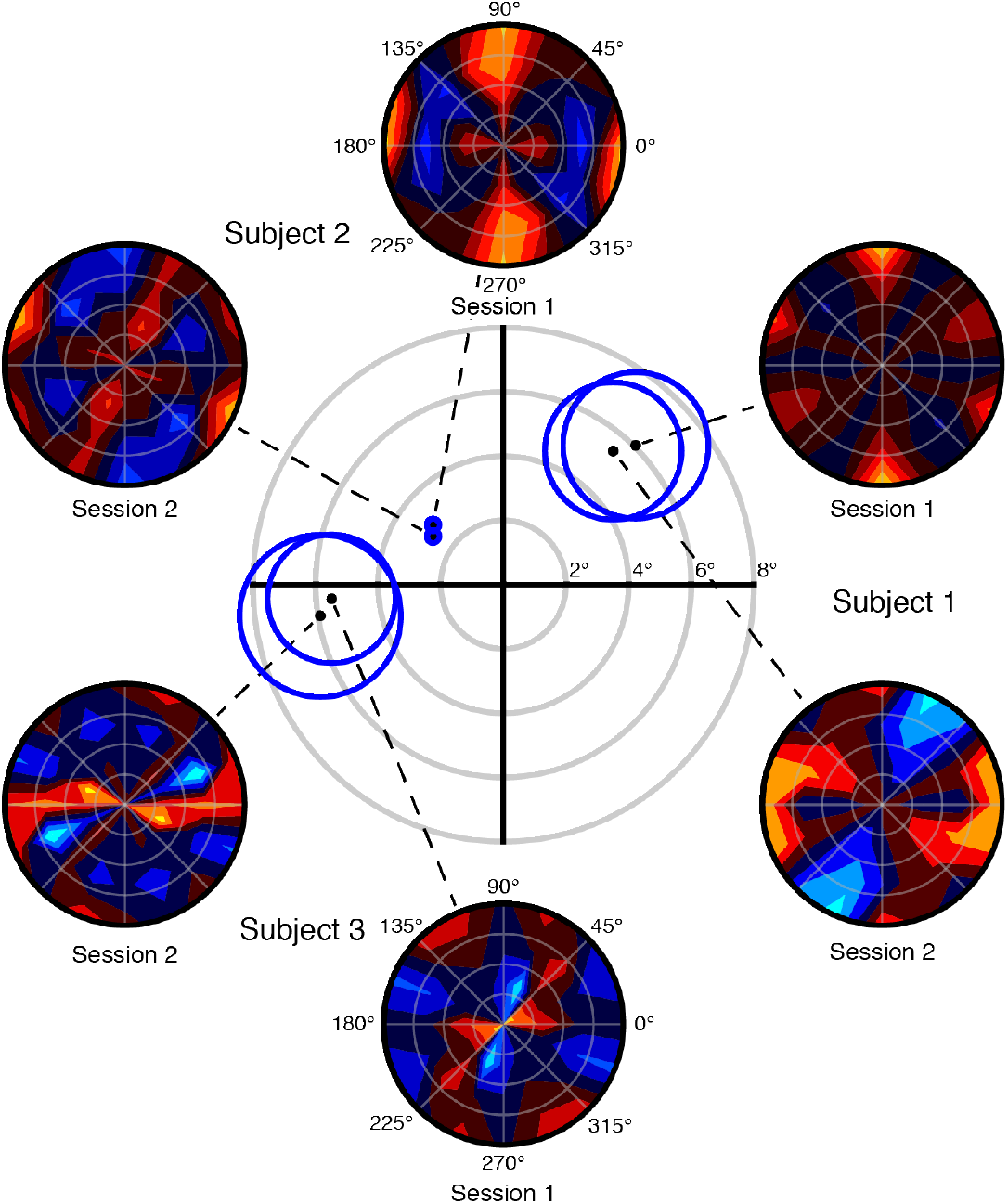
Consistency of response tuning across sessions. The retinotopic visual field location and orientation/spatial frequency tuning for each line is shown. The center plot represents the retinotopic visual field. For each line-scan session, a black dot represents the pRF center and a blue circle represents pRF size (2**σ** of Gaussian pRF model). The surrounding plots show orientation and spatial frequency tuning of each line. Orange and red are positive responses and blues are negative responses. The center of each plot represents high spatial frequencies and the outside represents low spatial frequencies.

### Response properties of cortical layers

Having established that we can identify cortical layers from quantitative mapping procedures, we asked whether layer-specific functional response properties could be discerned from line-scanning data. We computed pRFs and orientation/spatial frequency tuning on a voxel-wise basis. We found that pRF size is larger in infragranular (increasing 0.034° VA/100 μm from Layer 4 to white matter) and superficial layers (increasing 0.08° VA / 100 μm from Layer 4 to the cortical surface; Figure 5A), as has been previously described using electrophysiology in non-human primates (Gilbert, 1977; Hubel and Wiesel, 1977; Self et al., 2013; Snodderly and Gur, 1995) and using human fMRI (Fracasso et al., 2016). We computed the sharpness of orientation tuning for each voxel as the relative difference between preferred and perpendicular orientations. The sharpness of orientation tuning is higher in infragranular (0.17 a.u./100 μm) and supragranular layers (0.11 a.u./100 μm) compared to layer 4 (Figure 5B). This result corroborates electrophysiological observations (Hubel and Wiesel, 1977, 1959; Ringach et al., 2002; Self et al., 2019, 2013) and extends beyond the current fMRI literature.

**Figure 5.**
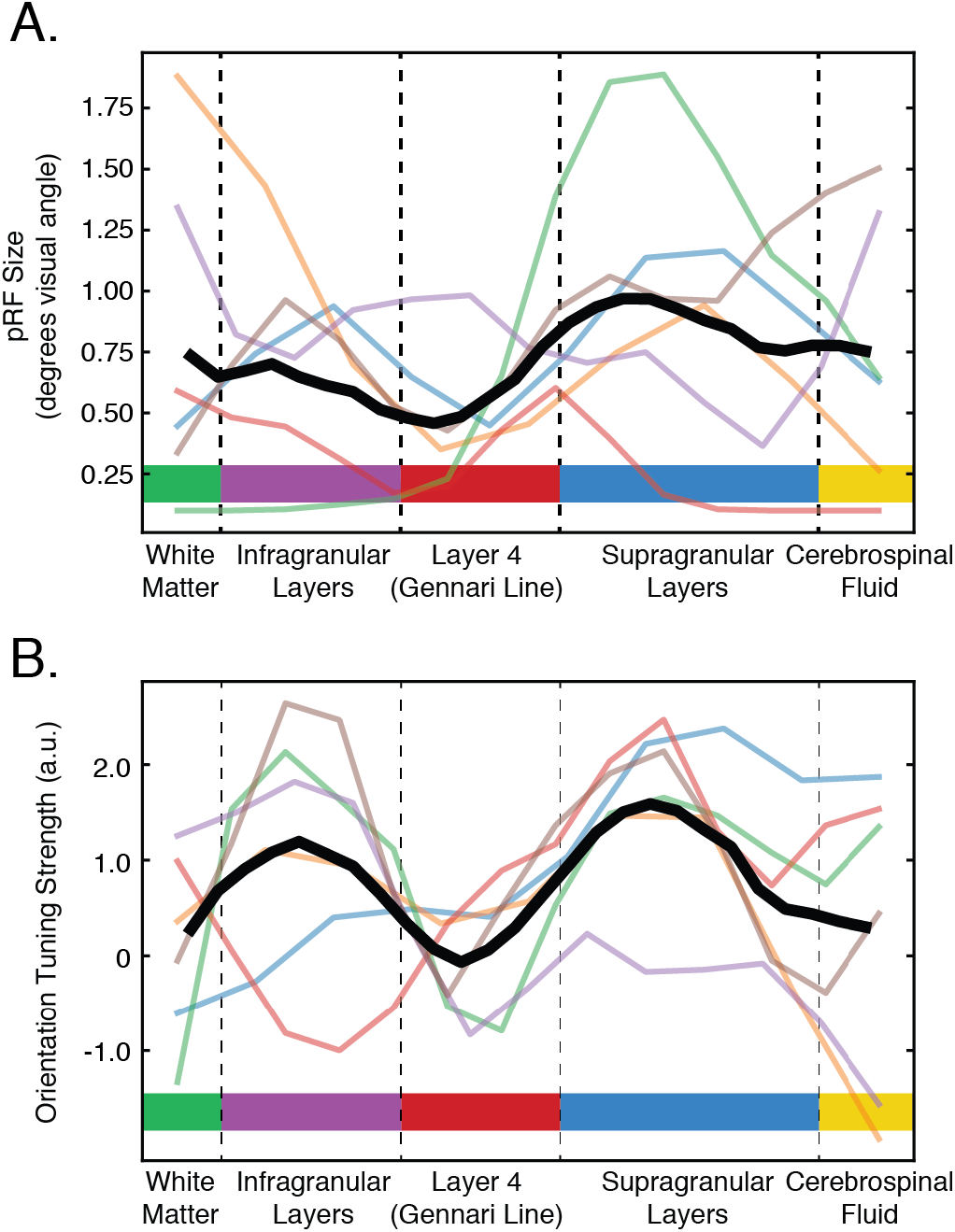
Layer-specific pRF tuning properties. A. pRF size is shown for each layer in degrees visual angle (**σ**). Individual subjects and sessions are shown and the black line shows the average across subjects, normalized for cortical thickness. B. Orientation tuning strength (defined as the relative difference between preferred and perpendicular orientations) across layers for individual subjects with the average shown in black.

## 4. Discussion

We have shown for the first time that line-scanning can be used to effectively record layer-specific functional signals from human cortex. Quantitative T_2_* and S_0_ mapping enabled us to identify cortical layers and functional recordings revealed layer-specific V1 tuning properties that had been previously observed in electrophysiological recordings in non-human primates. Consequently, we have demonstrated that line-scanning is a powerful non-invasive imaging technique for investigating mesoscopic functional circuits in human cortex.

With a spatial resolution of 200 μm, our functional recordings approach the resolution of individual cortical layers, which we used to compare tuning properties between Layer 4, infragranular layers and supragranular layers of human V1. Human layer fMRI studies must typically pool voxels spanning hundreds or even thousands of mm^2^ of cortex to differentiate responses in each of these layers (Polimeni et al., 2018). This poses a challenge to experiments probing responses of local neuronal circuits in vision (Hess and Field, 1999; Lamme, 1995; Roelfsema et al., 1998), as the receptive fields of large patches of cortex are not localized. However, our line-scan recordings have 10-12 voxels spanning cortical layers within a localized 3×3 mm^2^ patch of cortex. Line-scanning therefore has the potential to provide detailed descriptions of local circuit responses in human cortex.

Line-scanning fMRI can also help validate the correspondence of human neuroimaging data with what we know about neuronal and hemodynamic response coupling from animals. For instance, a recent study investigating BOLD responses in rat somatosensory cortex paired line-scanning with optogenetics (Albers et al., 2018) and found that optogenetic activation of specific excitatory neurons causally generated BOLD signals with an earlier onset compared to responses to sensory stimulation. This dissociation was explained by the fact that optogenetically-controlled neurons can be rapidly and precisely stimulated while sensory responses require intracortical transmission, effectively slowing hemodynamic responses. This study highlights the role of line-scanning fMRI to elucidate temporal dynamics of hemodynamic responses. Further, it represents a potential line of research in which advanced optogenetic control of neurons or neuronal compartments could be performed in rodents (as in Takahashi et al., 2016) and directly compared to human layer-specific responses to perceptual or cognitive functions.

Human fMRI has traditionally been regarded as a method lacking the temporal resolution required to detect dynamic functional changes we have access to with invasive animal recordings. For example, measuring oscillatory neuronal activity requires single or multi-unit recordings with millisecond precision. However, fluctuations in population responses can be captured in envelopes of power changes in electroencephalographic measures and average peristimulus histograms. Temporal resolutions of tens or hundreds of milliseconds are thus sufficient to describe dynamic activity between areas in functional networks or stimulus-evoked responses, respectively. In the current study, we credit our ability to read out detailed orientation tuning information from a relatively short fMRI experiment to an increased temporal resolution afforded by the line-scanning method. Natural stimuli such as the movies used in this study are functionally efficient for mapping tuning properties onto neuronal components because individual features fluctuate rapidly through time. These fast fluctuations are difficult to disentangle with the sampling rate of many fMRI studies (although see Ekman et al., 2017; Hennig et al., 2007; Lewis et al., 2016). However, the temporal resolution of our line-scanning recordings was sufficient to extract meaningful layer-specific responses, although limited by the physiological resolution of the hemodynamic response. We envisage the temporal resolution of human line-scanning as being a powerful tool to disentangle neuronal and hemodynamic input to human cortical layers by examining BOLD response onset times, as has been previously investigated in rodents (Silva and Koretsky, 2002; Tian et al., 2010; Yu et al., 2019).

While line-scanning offers a number of distinct advantages, limitations include an extremely limited field of view and susceptibility to motion. Regarding the limited field of view, we point out that the regions of interest in the current study included only 10-12 voxels each and they occupied a total volume approximately 20% smaller than a single voxel in a standard 3T fMRI study (21.6 mm^3^ compared to 27 mm^3^). The line-scanning method can therefore be described as distinctly tailored to the examination of mesoscopic processes.

Line-scanning is also susceptible to subject motion. In the current study, we were only able to perform motion correction in the high-resolution line direction (one degree of freedom). It is not possible to correct for motion in other directions, so a scan could be potentially lost if the subject moved more than 1-2 millimeters in either of those directions. It is therefore important to recruit highly experienced participants who move very little for line-scanning studies. However, in future studies this issue could be alleviated by performing prospective motion correction using an optical camera system (Mattern et al., 2019; Qin et al., 2009) or by incorporating motion navigators into line-scanning sequences (Hoinkiss and Porter, 2017).

Advancing innovations in cognitive neuroscience approaches is essential to driving breakthroughs in our understanding of the human brain. Animal models have provided functional, mechanistic and anatomical evidence across broad spatial and temporal scales, advancing theories and principles that guide our understanding of signals in human V1. Concurrently, the emergence of high-resolution, non-invasive brain imaging technologies has allowed us to measure human brain organization at the level of cortical layers and columns (Dumoulin et al., 2017; Lawrence et al., 2019). This is a landmark advance towards describing the multi-scale architectures and functional complexities of the human brain because these functional units of information processing reside at the mesoscopic scale (Larkum et al., 2018, 2009; Martino et al., 2017). High-resolution human brain data can therefore be strategically merged with data from animal models and simulations generating functional relationships and anatomical reconstructions across multiple levels of brain organization. As part of this, developing cutting edge non-invasive imaging approaches with increased resolution underpins our ability to map functional activity onto neural circuits, offering the prospect to ask transformative new questions of cognitive information processing.

## Acknowledgements

We would like to thank Sydney N. Williams and Laurentius Huber for discussions that led to improved line-scanning procedures in the current work. We would like to thank Jasper Fabius, Anna Makova and Alison Symon for assistance in acquisition. We would also like to thank Selena Morgan and Amaia Benitez for comments leading to improvement of the manuscript.

## Footnotes

1 This scan required second level (SAR)

